# Highly Multiplexed Fluorescence Microscopy with Spectrally Tunable Semiconducting Polymer Dots

**DOI:** 10.1101/2023.12.05.570289

**Authors:** Ziyu Guo, Chetan Poudel, Margaret C. Sarfatis, Jiangbo Yu, Madeline Wong, Daniel T. Chiu, Joshua C. Vaughan

**Affiliations:** Department of Chemistry, University of Washington; Seattle, WA, 98195, USA; Lamprogen, Inc.; Bothell, WA, 98021, USA; Department of Bioengineering, University of Washington; Seattle, WA, 98195, USA; Department of Physiology and Biophysics, University of Washington; Seattle, WA, 98195, USA

**Author notes:** These authors contributed equally to this manuscript (Ziyu Guo, Chetan Poudel).

## Abstract

Current studies of biological tissues require visualizing diverse cell types and molecular interactions, creating a growing need for versatile techniques to simultaneously probe numerous targets. Traditional multiplexed imaging is limited to around five targets at once. Emerging methods utilizing sequential rounds of staining, imaging, and signal removal can probe tens of targets but require specialized hardware, time-consuming workflows, and face some challenges with sample distortion and artifacts. Here we present a new method for highly-multiplexed fluorescence microscopy using semiconducting polymer dots (Pdots) in a single round of staining and imaging. Pdots are small, bright, and photostable fluorescent probes with a wide range of tunable Stokes shifts (20–450 nm). Multiple series of Pdots with varying excitation wavelengths allow for fast (<1 minute) and single-round imaging of up to 21 targets in brain and kidney. This method is based on a simple immunofluorescence workflow, efficient use of spectral space, standard hardware, and straightforward analysis, making it widely applicable for bioimaging laboratories.

## Introduction

Biological tissues are complex networks of cells that are arranged within intricate extracellular environments and that adapt during development, aging, and disease. Fluorescence microscopy is heavily used for the study of these complex specimens and particularly benefits from the high molecular specificity of immunofluorescence and related staining methods due to their powerful ability to quantify the distributions of multiple molecules of interest within the specimen. Commonly used fluorophores in microscopy have absorption and emission spectral bandwidths of 30–60 nm and a Stokes shift of 20–30 nm between the absorption and emission maxima. Because each fluorophore occupies a combined ∼70nm of spectral bandwidth, the number of fluorophores concurrently visualized across the visible and near-infrared (400–750 nm) is commonly limited to ∼5.

Prior work has sought to increase fluorescence multiplexing capabilities using various immunolabeling strategies and detection techniques (*1*). One family of techniques uses sequential cycles of staining and imaging and avoids spectral overlap by chemically inactivating or removing fluorescent dyes after each cycle (e.g., MxIF (*2*), t-CyCIF (*3*), IBEX (*4*), and SWITCH (*5*)). A second family of sequential multiplexing techniques uses a single step to immunostain tissues with DNA-barcoded antibodies, followed by a sequential readout of barcodes using complementary fluorescent oligonucleotides (e.g., CODEX (*6*) and Immuno-SABER (*7*)). Multiplexed transcriptomics methods are a third family of techniques that map mRNA distributions across cells and tissues (e.g., MERFISH (*8*) and seqFISH (*9*)). These methods also use a single round of staining with barcoded probes for gene panels, followed by sequential rounds of reagent delivery and imaging to read out the barcodes and spatially localize the transcripts. Despite the large number of targets that can be probed, all of these sequential approaches encounter major limitations: extended times for probe delivery and incubation (usually 20–120 minutes or longer per cycle and on order of a day or more per experiment) that restrict the application to imaging of thin tissues; the potential for tissue distortion between rounds and for unsuccessful inactivation of probes; complex and computationally intensive probe design and analysis; and the requirement for specialized and automated fluidic handling equipment and dedicated imaging instruments.

Instead of multiple rounds of staining and imaging, a single round of highly multiplexed imaging would be desirable in many circumstances. Laser or ion beam ablation followed by mass spectroscopy can achieve higher plex imaging in a single round (*10, 11*), but this approach involves a tedious workflow with low throughput (e.g., pump-down of the sample chamber), is destructive, and requires expensive instrumentation. A fluorescence-based technique would alleviate some of these requirements, in principle, although commonly used microscopy fluorophores have a small Stokes shift of 20–30 nm that, together with the 30–60 nm excitation and emission bandwidths, limits the number of fluorophores that may be used. In contrast, recently developed spectrally tunable semiconducting polymer dots (Pdots) (*12*–*23*) exhibit a variable Stokes shift of 20–450 nm which can be tuned by varying the composition of the emitting Pdots (*17, 18, 20, 21*). This tunability of Stokes shift in Pdots provides much more spectral real estate for multiplexing than is possible using conventional fluorescent dyes. The related field of flow cytometry has led the way in pushing the limits of fluorescence multiplexing. However, many of the long Stokes-shift fluorophores that made high multiplex flow cytometry possible, such as phycoerythrin (PE) tandem dyes and the BD Brilliant series of polymer dyes, can be prone to photobleaching and are thus unsuitable for imaging. In contrast, Pdots are photostable and can have similar or better photostability as water-soluble Qdots (quantum dots) (*12*–*14, 20*). (A comparison of the photostability of Pdots and BD Brilliant dyes is shown in fig. S1.) Pdots are also brighter than small-molecule organic dyes or water-soluble Qdots due to their larger absorption cross-section (*14, 16, 17, 23*).

Here we demonstrate highly multiplexed imaging of up to 21 targets in a single round of immunostaining and imaging by using only three series of Pdots (a Pdot series is defined by its excitation peak, such as a 355 nm excitable series of Pdots comprising over 10 Pdots - each with a distinct emission peak from below 400 nm to over 800 nm). With computational processing using linear unmixing (*24*) after a single round of image acquisition, overlapping spectral signals of multiple Pdots can be unmixed and reassigned to their corresponding channels to provide reliable signal separation and distinct multi-target imaging. We demonstrate the versatility of our technique by using both direct and indirect immunolabeling of tissues (mouse kidney and brain), and by making both separate and concurrent use of excitation- and emission-multiplexed imaging.

## Results

### Semiconducting Pdots with Spectrally Tunable Properties

Pdots are semiconducting nanoparticles prepared by collapsing hydrophobic semiconducting polymers into a compact “dot”, the size of which can be tuned from ∼5 nm to over ∼20 nm (*12, 13, 15, 19*). The brightness of Pdots is generally very high, owing to their large absorption cross-section. In fact, by incorporating a Qdot into a Pdot, the brightness of the Qdot can be increased by up to an order of magnitude in comparison with a water-soluble Qdot, alone (*16*). The photophysical property of Pdots that is of particular relevance to the current study is amplified energy transfer, which enables the Stokes shift to be tuned from ∼20 nm to over 400 nm (*20, 21*). This characteristic, together with narrow emission bands, allows for the design and creation of a large color panel (e.g., 10) for a single wavelength excitation. Unlike Qdots, the absorption peak of Pdots can also be tuned to enable excitation multiplexing. Combining excitation and emission multiplexing enables the creation of a very large color panel that numbers in the many tens. These favorable properties of Pdots have now facilitated highly multiplexed flow-cytometry analysis of single cells, and Pdots are commercially available as the StarBright series from Bio-Rad, with over 30 colors launched thus far. Unlike many of the tandem dyes used traditionally in flow cytometry, most Pdots possess good photostability and may enable highly multiplexed fluorescence imaging. Additionally, Pdots can be designed with further improved photostability or used under conditions that improve their photostability (*22*).

For multiplexed imaging in this study, we used three series of spectrally distinguishable Pdots with absorption maxima near 355 nm, 405 nm, or 488 nm for optimal excitation with commonly available laser lines. For each series, we used 6–8 Pdots with a range of emissions covering the visible and near-infrared spectrum, from below 400 to over 800 nm. The excitation and emission spectra of this palette of 22 Pdots are shown in Fig. 1.

**Fig. 1.**
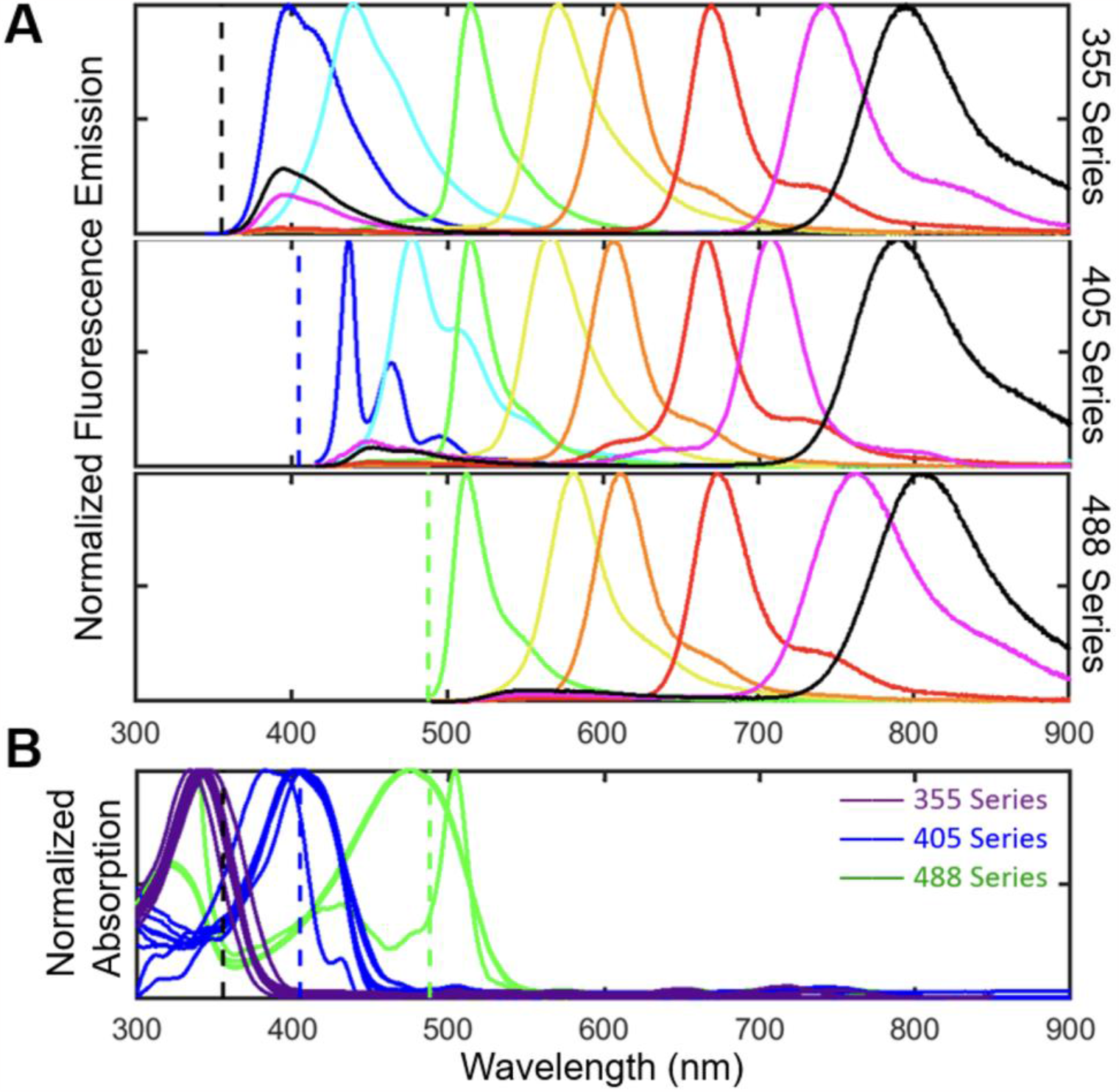
Emission (A) and excitation (B) spectra of three series of Pdots. Each series contains 6–8 Pdots with emission wavelengths that span the visible and near-infrared spectrum from below 400 to over 800 nm. Dashed lines indicate the excitation wavelengths used (365, 405, and 488 nm).

In order to measure fluorescence from about 400 to 800 nm for our panels of Pdots, we sequentially recorded images using a set of bandpass emission filters (fig. S2) for a given excitation wavelength, then repeated the process for up to three excitation wavelengths. While the emission filters were selected to match each Pdot, some emission bleedthrough was present due to spectral overlap of the Pdots within each Pdot series (Fig. 1A). This emission bleedthrough is most notable for Pdots in adjacent spectral bands and for Pdots with a principal emission peak of 700–800 nm that exhibit a minor emission peak with a small Stokes shift of 50–100 nm. In addition, the excitation spectra of the three Pdot series partly overlap at the excitation wavelengths, leading to some cross-excitation (Fig. 1B). For instance, at 365 nm excitation, 355ex-Pdots (the 355 nm excitation series of Pdots) are excited at ∼80% of their excitation maximum while the 405ex-Pdots and 488ex-Pdots are also cross-excited at ∼20–30% of their excitation maximum. To address emission-bleedthrough and cross-excitation, we employed a linear unmixing algorithm in MATLAB to reassign signals to their corresponding channels based on calibration data measured for individual Pdots. This procedure allowed us to generate unmixed images that correct for crosstalk.

### Emission multiplexing: Visualizing eight targets using a single excitation wavelength in a single round

To concurrently label and visualize eight key structures in mouse brain using emission multiplexing with Pdots excited at 405 nm, we first selected and validated seven primary antibodies and the lectin wheat germ agglutinin (WGA) with distinct staining patterns in mouse brain. The panel of seven primary antibodies was selected to also be antigenically distinct (i.e., mouse IgG1, mouse IgG2a, mouse IgG2b, mouse IgG3, rabbit, rat, and chicken) so that they could each be uniquely targeted by specific secondary antibodies (i.e., goat anti-mouse IgG1, goat anti-mouse IgG2a, goat anti-mouse IgG2b, goat anti-mouse IgG3, donkey anti-rabbit, donkey anti-rat, and donkey anti-chicken; see table S1 for details). Next, eight Pdots from the 405 nm series with emission peaks ranging from 430 nm to 800 nm were conjugated to secondary antibodies or directly to WGA. Fig. 2 shows a 100 µm thick brain tissue section stained for eight targets using these probes and then imaged within a few seconds in a single round of automated imaging using matched filter sets. The images displayed in Fig. 2 were linearly unmixed from raw data to correctly reassign bleedthrough from each spectral band to the corresponding stain; see the methods for details on the linear unmixing procedure and fig. S3 for a comparison of the raw and unmixed images. The secondary antibody-Pdot conjugates bind to primary antibodies accurately with a low background signal. Anti-MBP labeled oligodendrocytes, anti-GFAP labeled astrocytes/glial cells, WGA labeled N-Acetylglucosamine, anti-NeuN labeled neuronal nuclei, anti-NFH labeled heavy neurofilaments, anti-AQP4 labeled aquaporin-4 water channel protein, and anti-Tuj1 labeled beta-tubulin III on neural stem cells.

**Fig. 2.**
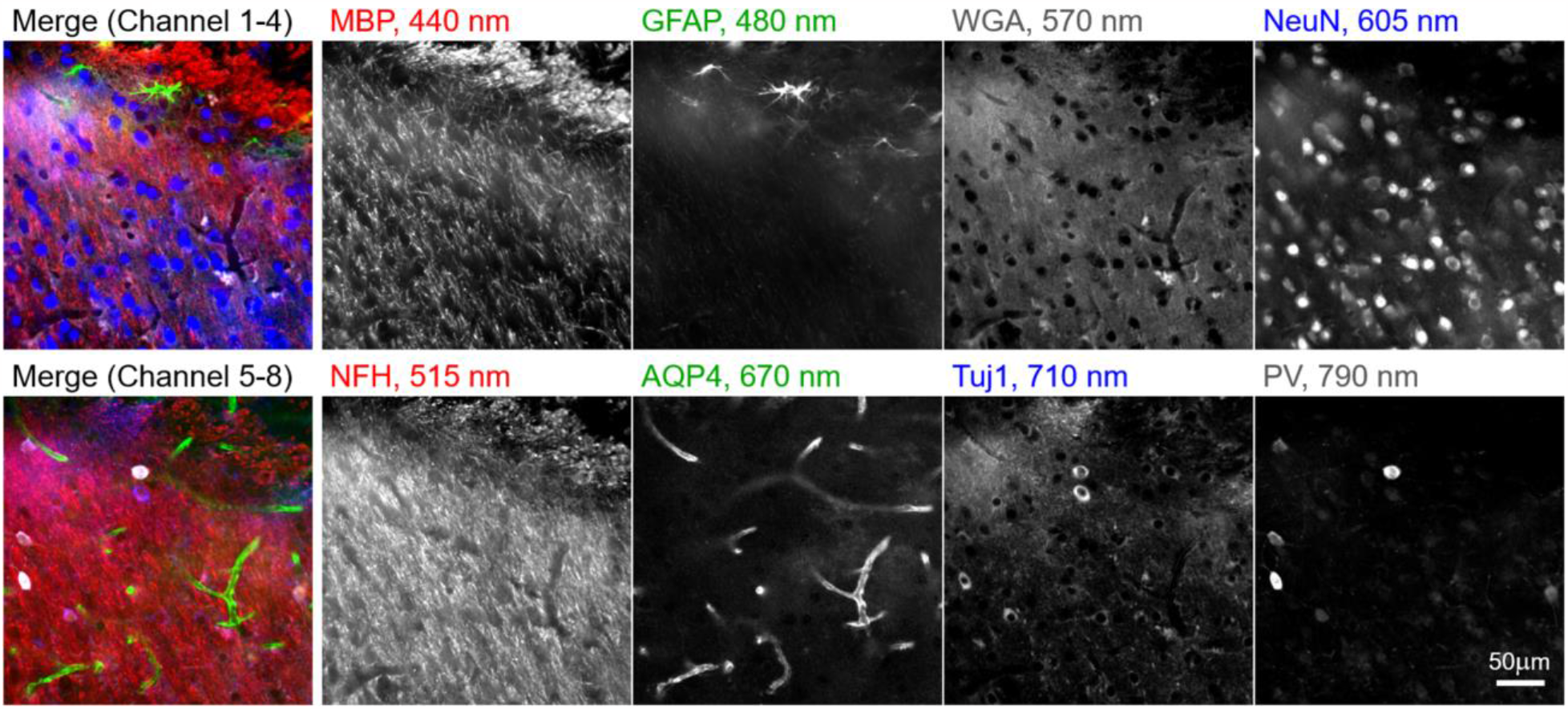
Emission multiplexing with the 405ex-Pdot series: A 100 μm thick brain slice was stained with eight different Pdots conjugated to a set of seven secondary antibodies and the lectin wheat germ agglutinin. The specimen was illuminated with 405 nm light on a widefield microscope and the signal measured through different bandpass filters was linearly unmixed. (MBP = myelin basic protein, GFAP = glial fibrillary acidic protein, WGA = wheat germ agglutinin, NeuN = neuronal nuclei, NFH = neurofilament H, AQP4 = aquaporin-4, Tuj1 = tubulin beta III, PV = parvalbumin.)

In a related experiment for 8-channel emission multiplexing with the 405 nm Pdot series, we used a set of five secondary antibody-Pdot conjugates (anti-mouse IgG1, anti-mouse IgG2a, anti-mouse IgG2b, anti-rabbit, anti-rat), a direct antibody-Pdot conjugate, and two Pdot-lectin conjugates, to label and image a mouse kidney tissue section (fig. S4). We also performed 7-channel emission multiplexing with the 355ex-Pdot series in mouse brain and mouse kidney using a mixture of Pdot-secondary antibody conjugates, Pdot-primary conjugates, and Pdot-lectin conjugates (figs. S5 and S6). Together, this collection of experiments demonstrates the feasibility of using emission multiplexing with Pdots to achieve up to 8-channel imaging with a single excitation wavelength.

### Excitation multiplexing: Visualizing three targets using a single emission filter in a single round

In addition to emission multiplexing, in which one excitation wavelength excites a set of Pdots that fluoresce across a range of emission wavelengths, Pdots can also be used for excitation multiplexing, in which Pdots that emit at the same wavelength are selectively excited using different illumination wavelengths. For example, in Fig. 3, we demonstrate three-channel excitation-multiplexed imaging of mouse brain tissue using Pdots that emit at 670 nm and of mouse kidney tissue using Pdots that emit at 800 nm. In each case, the Pdots are selectively excited via illumination at 365 nm, 405 nm, and 488 nm. Unlike emission multiplexing, in which bleedthrough into neighboring emission channels is observed, here we encounter some cross-excitation of Pdots that produces channel crosstalk (e.g., 405 nm illumination undesirably cross-excites the 488ex-Pdots ∼20%, see Fig. 1B). We corrected this cross-excitation in a similar fashion as before using linear unmixing (see fig. S7 for a comparison of the raw and unmixed data). This ability to perform excitation multiplexing is unavailable to Qdots because they do not have distinct absorption peaks.

**Fig. 3.**
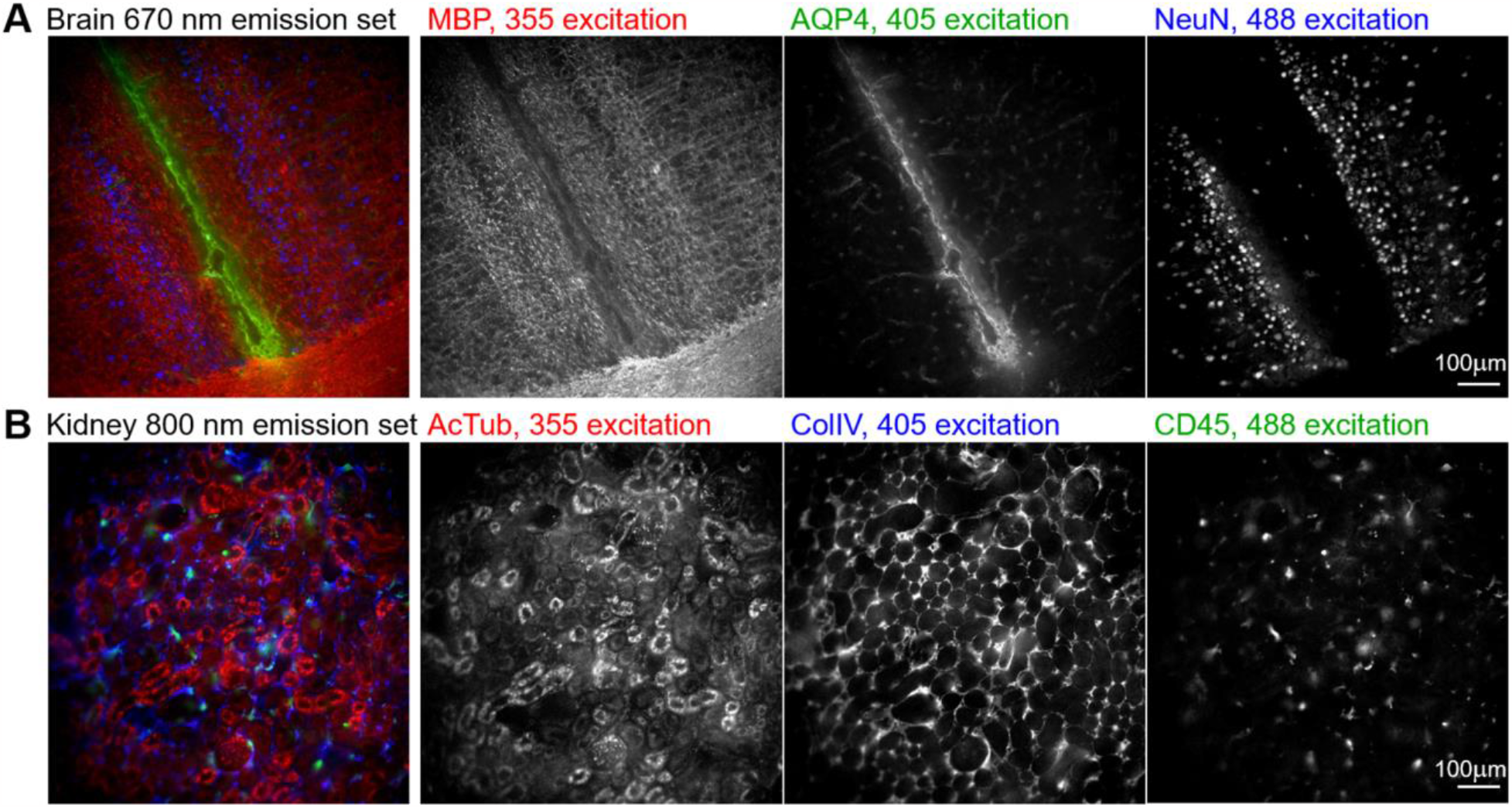
Excitation multiplexing: Three-channel excitation-multiplexed imaging of 100 µm thick tissue sections of (**A**) mouse brain and (**B**) mouse kidney. Each set of three Pdots was illuminated at 365 nm, 405 nm, and 488 nm; a 667/30nm emission filter was used for brain, and a 800/60nm emission filter for kidney. (MBP = myelin basic protein, AQP4 = aquaporin-4, NeuN = neuronal nuclei, AcTub = acetylated tubulin, ColIV = collagen 4, CD45 = leukocyte common antigen.)

### Excitation and emission multiplexing: up to 21-plex imaging in a single round

Having separately demonstrated the possibilities of using Pdots for excitation- and emission-multiplexed imaging, we sought to combine them using our three series of Pdots to stain and visualize up to 21 targets in a single round. Using indirect immunofluorescence to stain a large number of targets in a single round is impractical due to limitations in the number of independent primary/secondary antibody combinations. Therefore, when creating larger palettes of probes, we chose to conjugate the Pdots directly to primary antibodies or lectins (see table S3). We validated independent antibodies or lectins as labeling their respective targets of interest, conjugated each antibody or lectin to a Pdot, and performed careful crosstalk calibration measurements for each Pdot. Using this procedure, we were able to perform 14-plex imaging of a mouse brain section and 21-plex imaging of a mouse kidney section, with the data acquisition for each being performed in under a minute (Fig. 4–5). As before, details of filters, antibodies, and Pdot conjugations are listed in tables S1–S3, and the linear unmixing procedure is described in the methods section.

**Fig. 4.**
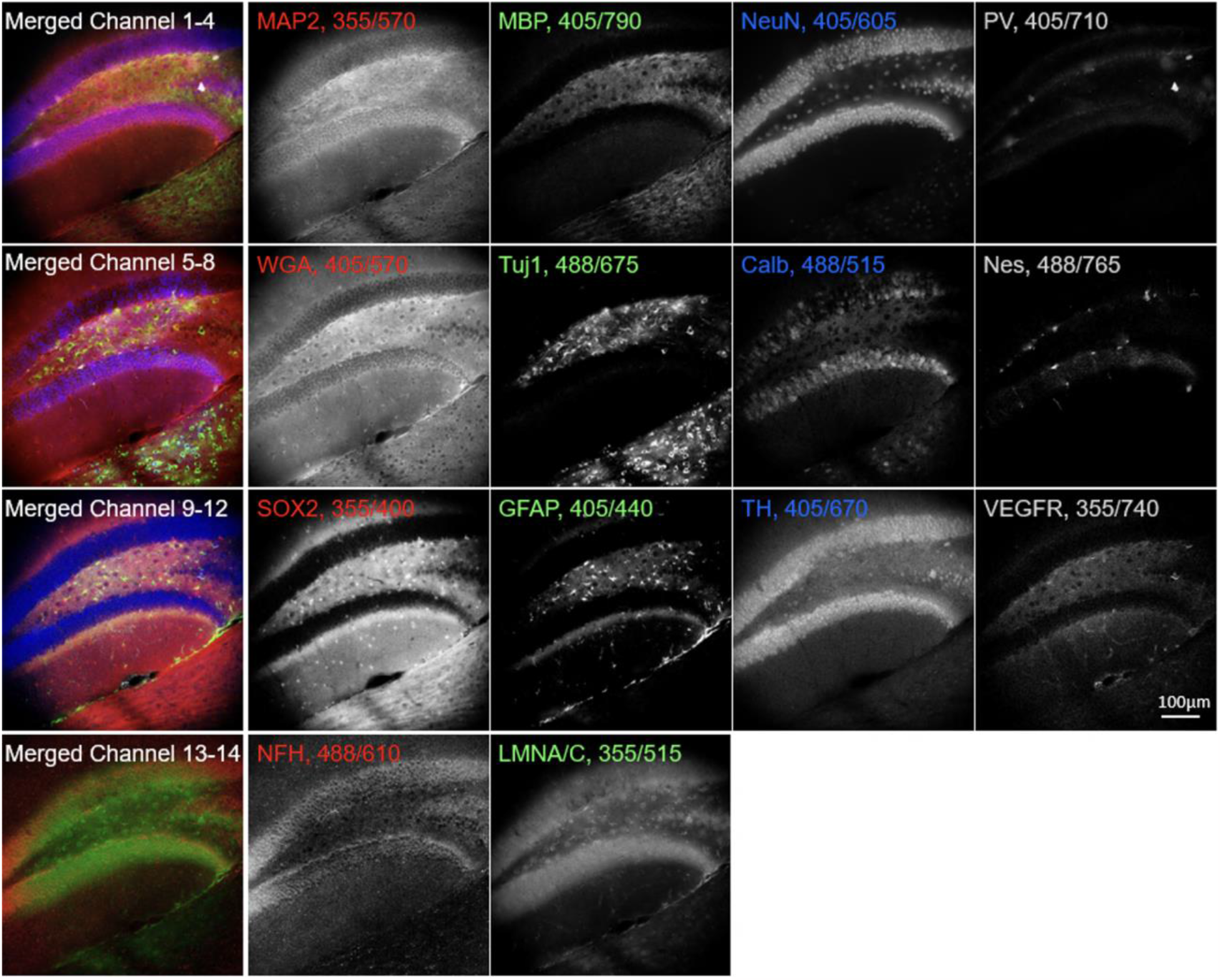
Excitation and emission multiplexing in brain tissue: A 50 µm thick mouse brain section stained with 14 Pdots conjugated to primary antibodies and lectins. The tissue was imaged using a combination of three excitation wavelengths and eleven emission filters. The left column shows four partial 4-channel composite images; the four columns to the right show corresponding grayscale images. Labels indicate the protein target or lectin (WGA) together with Pdot excitation/emission wavelengths. (MAP2 = microtubule associated protein 2, MBP = myelin basic protein, NeuN = neuronal nuclei, PV = parvalbumin, WGA = wheat germ agglutinin, Tuj1 = tubulin beta III, Calb = calbindin, Nes = neuroepithelial stem cell protein, SOX2 = sex determining region Y-box 2, GFAP = glial fibrillary acidic protein, TH = tyrosine hydroxylase, VEGFR = vascular endothelial growth factor receptor 3, NFH = neurofilament H, LMNA/C = lamin A/C.)

**Fig. 5.**
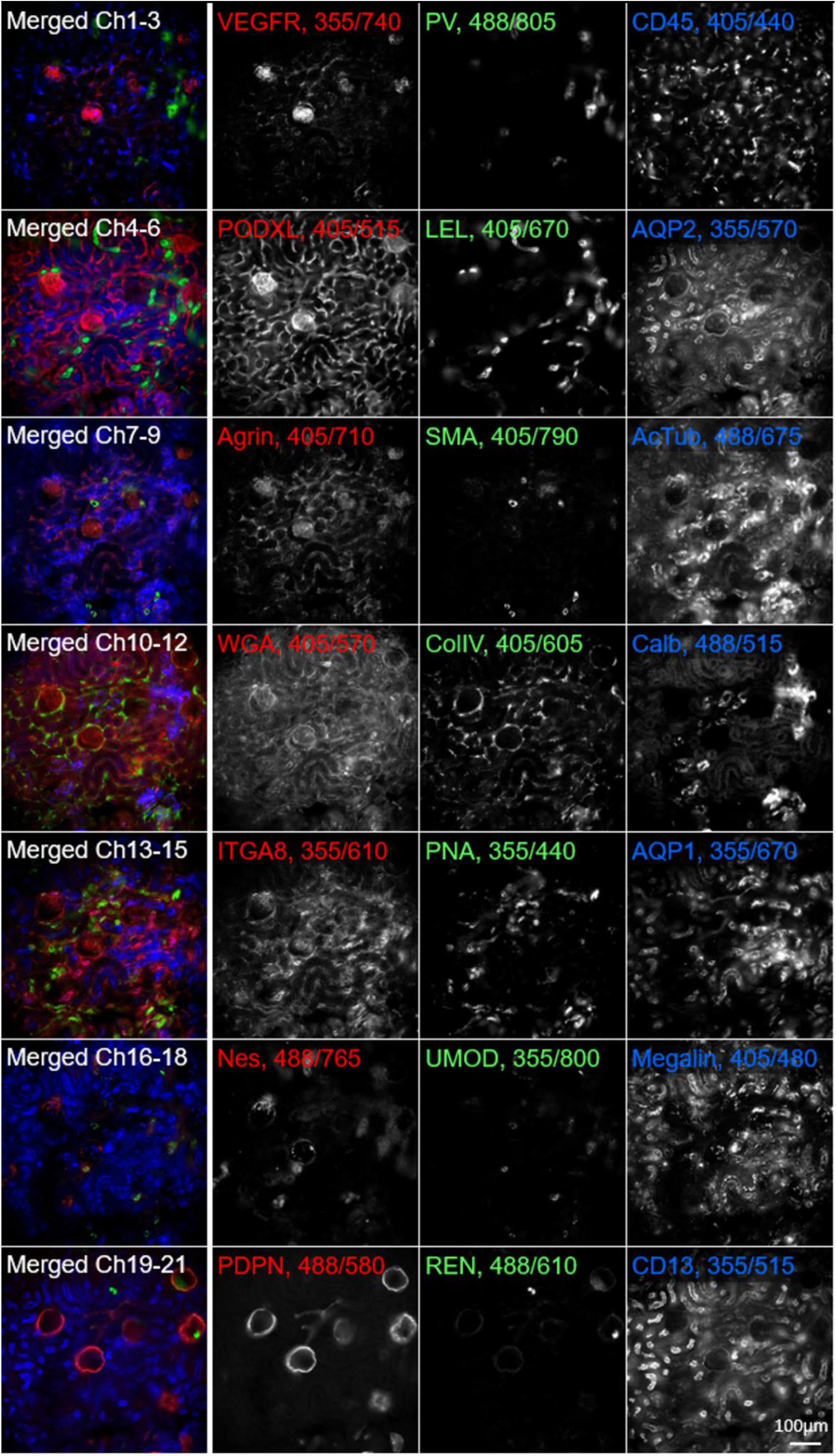
Excitation and emission multiplexing in kidney tissue: A 50 µm thick mouse kidney section stained with 21 Pdots conjugated to primary antibodies and lectins. The tissue was imaged using a combination of three excitation wavelengths and eleven emission filters. The left column shows seven 3-channel composite images of the same tissue region; the three columns to the right show corresponding grayscale images. Labels indicate the protein targets or lectins (LEL, WGA, PNA) together with Pdot excitation/emission wavelengths. (VEGFR = vascular endothelial growth factor receptor 3, PV = parvalbumin, CD45 = leukocyte common antigen, PODXL = podocalyxin, LEL = lycopersicon esculentum tomato lectin, AQP2 = aquaporin 2, SMA = alpha-smooth muscle actin, AcTub = acetylated tubulin, WGA = wheat germ agglutinin, ColIV = collagen 4, Calb = calbindin, ITGA8 = integrin alpha 8, PNA = peanut agglutinin, AQP1 = aquaporin 1, Nes = neuroepithelial stem cell protein, UMOD = uromodulin, Megalin = low density lipoprotein-related protein 2, PDPN = podoplanin, REN = renin, CD13 = platelet endothelial cell adhesion molecule-1.)

## Discussion

Multiplexed imaging is increasingly important to probe the interactions of various cell types in disease and in the context of discovery efforts such as tissue atlases. Our technique provides broad applicability since Pdots can be bound to commercially available conventional antibodies, which offers flexibility in the choice of cellular markers, antibody-Pdot combinations, etc. Compared to iterative staining methods (*2*–*7*), our method is simpler and faster, and does not require probe inactivation reagents or suffer from tissue distortion. We make use of standard imaging hardware available in most bioimaging labs, unlike other multiplexing approaches that use more sophisticated hardware or may need dedicated instruments for automated fluidics or multi-day acquisitions (*6*–*11*). We have demonstrated that in a single round of highly multiplexed imaging, we can concurrently investigate up to 21 targets on a single 50 µm thick tissue specimen using three series of Pdots with excitation at 365 nm, 405 nm, and 488 nm. Additional 560 nm and 647 nm series of Pdots would enable excitation multiplexing (*25*) to a larger extent than we demonstrated here, and the number of targets could be increased further to ∼30 in a single round through a combination of excitation and emission multiplexing with a 5 series of Pdots. Combining this approach with antibody stripping or Pdot inactivation (e.g. via quenching or bleaching) may make it possible to measure over 100 targets in only 4–5 rounds of imaging.

In contrast to the above benefits, the use of Pdots for highly multiplexed imaging does face some challenges. The process of conjugating antibodies to Pdots places some restrictions on the choice of antibodies and requires careful pre- and post-validation of antibody performance. For this study, we tested 53 Pdot-primary antibody (or Pdot-lectin) conjugates (table S3). Of these 53 conjugates, 31 consistently produced strong staining of the correct molecular targets, 9 produced weak staining, and 13 showed no staining or unsuitably high background indicative of non-specific binding. These results show an estimated ∼60% success rate for Pdot conjugation. The failure rate of Pdot-primary antibody conjugates (∼40% not binding their targets efficiently) is similar to that of DNA-conjugated primary antibodies and other approaches involving custom primary conjugates to conventional dyes (*3*). Purchasing different primary antibodies against the same target and repeating the conjugation often led to identifying a successful conjugate.

As part of our image processing workflow, we used linear unmixing - a simple and popular technique in the field of multiplexed imaging which uses raw/mixed image data and a reference/calibration matrix to generate unmixed spectral images (*24, 26*). Despite the technique’s simplicity, it requires a pre-measurement of all the Pdots individually in each imaging channel before the unmixing can be performed. However, this calibration can be performed once for all subsequent experiments where the same imaging settings are used. Techniques like hyper-spectral phasors (HySP) and blind source separation methods can unmix signals without prior measurements of reference spectra but generally perform worse when there are larger numbers of mixed channels (*27*–*30*). For instance, PICASSO (*30*) uses a sophisticated unmixing algorithm that seeks to maximize the differences between multiple channels but has some potential limitations in cases where channels may be innately similar to one another.

We also noticed higher levels of background when illuminating the 355 nm series of Pdots; at these shorter excitation wavelengths, biological tissues exhibit strong autofluorescence that can mask the specific staining or impact the unmixing calculation, generating images with autofluorescence contamination in the short wavelength channels. The use of thinner tissue sections or incorporating corrections for autofluorescence may improve the multiplexing performance of these Pdots. While most Pdots are bright and photostable, we noticed that some Pdots with larger Stokes shifts exhibit blue-shifting of their spectrum in a power density dependent manner when exposed to bright focused light (e.g., when using a confocal microscope with a high-NA objective lens). This spectral shift can introduce errors during linear unmixing, as signals tend to be weaker for Pdots with larger Stokes shift (i.e., redder emission channels). This may limit the current applications of Pdot multiplexing to microscopies that use lower power densities, such as widefield and lightsheet microscopy. In the future, engineering Pdots that do not exhibit spectral shifts would make them more broadly applicable and better suited to microscopy modes that involve very high excitation power densities.

With the availability of additional sets of Pdots excitable at 561 nm and 640 nm, together with increasing emission multiplexing and better compensation methods, it will be possible to further increase the number of targets that can be studied in a single round of staining and imaging. While fewer emission bands can be accommodated for these redder excitation wavelengths, incorporating these new Pdot series could allow for probing ∼30 targets in a single round. We are also investigating approaches to engineer these Pdots to have narrower excitation and emission bands to reduce bleedthrough and cross-excitation, which will further increase the number of targets that can be probed in a single round of imaging.

## Materials and Methods

### Experimental Design

This study aims to develop a new approach to highly-multiplexed fluorescence microscopy with a simple workflow and wide applicability in bioimaging laboratories with standard imaging hardware. The experimental design is based on the use of novel fluorescent semiconducting nanoparticles (Pdots) that have a large range of Stokes shifts and are spectrally distinct from each other, thus enabling combinations of excitation and emission multiplexing. Pdots are conjugated to commercially available antibodies of interest. By designing all experiments to be conducted in a single round of staining and imaging, our approach emphasizes simplicity, speed, and applicability to routine microscopy procedures.

### Absorption and emission spectra

Semiconducting Pdots were diluted 1:100 in PBS-azide (phosphate buffered saline containing 3 mM sodium azide) and measured using a PerkinElmer LS55 fluorometer. Three excitation wavelengths—355 nm, 405 nm, and 488 nm—were used to generate emission by the corresponding Pdot series.

### Mouse brain and kidney sample preparation

All procedures used in this study were approved and performed under the guidelines from the University of Washington Animal Use Training Session and the regulations from Institutional Animal Care and Use Committee. Three mice were used in this study: Mouse 1, a 6-month-old C57BL/6 Cre-ERT2 female; Mouse 2, a 2-month-old C57BL/6 wildtype female; and Mouse 3, a 5.5-month-old B6*P14 transgenic female. Tissues from these mice did not exhibit any fluorescent reporters. Immediately after CO_2_ euthanasia, mice were cardiac-perfused with PBS-azide and 4% paraformaldehyde (PFA), each for 5 min. The kidneys, with renal capsule removed, were harvested and fixed in 10 mL of 4% PFA for 1 h. The brain was fixed overnight in PFA. After fixation, the brain was embedded in agarose gel and sliced into 50 µm and 100 µm thick sections on a vibratome along the coronal plane. The kidneys were sliced without embedding into 50 µm and 100 µm sections along the coronal plane. All sections were stored in PBS-azide at 4 ºC until further use.

### Screening and validation of antibodies

The full list of primary and secondary antibodies used in this study is provided in table S2– S3. We validated the performance of primary antibodies by staining tissues with the primary antibody and then a pre-validated secondary antibody conjugated to conventional fluorescent dyes (such as ATTO 488, Alexa Fluor 568, or ATTO 647N). We confirmed that the antibodies bound the correct targets in desired tissues by comparing our images with the literature. Secondary antibodies were validated in a similar manner but in combination with a pre-validated primary antibody. After this first round of validation, the antibodies (primary or secondary) were conjugated to Pdots. In some cases, the conjugation could change the antibody’s structure, antigenicity, or binding efficiency. We therefore re-validated each antibody after conjugation to Pdots to confirm target binding via imaging. The process of conjugating antibodies to Pdots places some restrictions on the choice of antibodies. Some components typically found in the antibody buffer solution, such as bovine serum albumin, gelatin, albumin, and Tris buffer, can interfere with the conjugation reaction and make the yield harder to quantify. We chose to use antibodies without these components, which sometimes limited us to using antibodies that were not used widely in other papers.

### Conjugation of antibodies to Pdots

The preparation of Pdot-conjugates has been described in detail in prior publications (*12*– *14, 17, 21*). Briefly, Pdots were conjugated to antibodies or lectin via a 1-ethyl-3-(-3-dimethylaminopropyl) carbodiimide hydrochloride (EDC) catalyzed reaction, in which 0.25 mL of Pdot solution was mixed with 3 uL of EDC and 60 uL of an antibody solution, and reacted for 3 h. The final solution was then loaded onto a gel filtration column to purify the Pdot-antibodies from free unreacted antibodies.

### Immunostaining

Tissues were incubated in block/permeabilization solution (3% bovine serum albumin and 0.5% Triton-X 100) for at least 4 h at room temperature on a shaker. All tissues were stained in a single round. For indirect immunofluorescence, tissues were stained overnight with primary antibody and overnight with Pdot-conjugated secondary antibody. For direct immunofluorescence, tissues were stained for 24 h. Tissues were washed with PBS-azide 3 times for 10 min each and were imaged in a single round with the proper filter combinations (table S1). Luminescence tends to be dimmer for Pdots with larger Stokes shifts and the quantum efficiency of standard sCMOS cameras also declines towards the near-infrared region. To counter these issues, we paired the brightest immunostains with the dimmest Pdots and increased the concentration of the dimmest stains to achieve similar levels of signal intensity for each channel.

### Imaging

All imaging was performed on a Nikon Ti2 Eclipse microscope equipped with an automated objective z-drive for focusing, automated XY stage (Applied Scientific Instrumentation), motorized filter cube turrets, an assembly of motorized filter wheels (Thorlabs), and a scientific CMOS camera (Hamamatsu Orca Flash 4.0v3). Excitation was achieved by filtering the white light output of a SOLA U-nIR light engine (Lumencor) using three excitation filters. All dichroic, excitation, and emission filters are listed in table S1 and were purchased from Semrock or Chroma. Emission filters were optimized to capture a band of 20–60 nm at the emission peaks of each Pdot.

### Linear unmixing

Mixed signals due to spectral overlap in each channel were computationally unmixed using a custom analysis script in MATLAB. The emission profiles of each conjugated Pdot in solution were measured across the available emission filter channels to record the proportion of crosstalk and signal bleedthrough to neighboring channels. During initial stages of the project when excitation and emission multiplexing were performed separately, we used a simpler calibration procedure for just the few necessary channels. As more Pdots were conjugated and the full palette of 22 distinct Pdots was available, we performed a full calibration for each pure Pdot in solution (i.e., using each combination of illumination wavelength and emission channel for each Pdot) to produce a 22 × 22 calibration matrix. The inverse of the complex conjugate of this calibration matrix was multiplied to the raw mixed datasets to generate unmixed datasets using non-negative least squares fitting on a pixel-by-pixel basis.

### Statistical Analysis

Image data was processed using established software tools (e.g., ImageJ and MATLAB). Quantitative measurements, such as signal intensities, were extracted from regions of interest. Raw data, including images and measurement values, are available in supplementary data.

## Acknowledgments

We gratefully acknowledge assistance from Dr. Chenyi Mao in the initial stages of this project.

## Author contributions

Conceptualization: ZG, CP, JCV, DTC Methodology: ZG, CP, MCS, JY, MW, DTC, JCV

Investigation: ZG, CP Visualization: ZG Supervision: DTC, JCV

Writing—original draft: ZG, CP, JCV, DTC

Writing—review & editing: ZG, CP, MCS, JY, MW, DTC, JCV

## Data and materials availability

All relevant data needed to evaluate the conclusions in the paper are present in the paper and/or the Supplementary Materials. Please contact the authors to request any additional data or the MATLAB source code used for linear unmixing of mixed signals.

