## Supplementary Materials for "Highly Multiplexed Fluorescence Microscopy with Spectrally Tunable Semiconducting Polymer Dots"

**This PDF file includes:**

Figs. S1 to S7
Tables S1 to S4

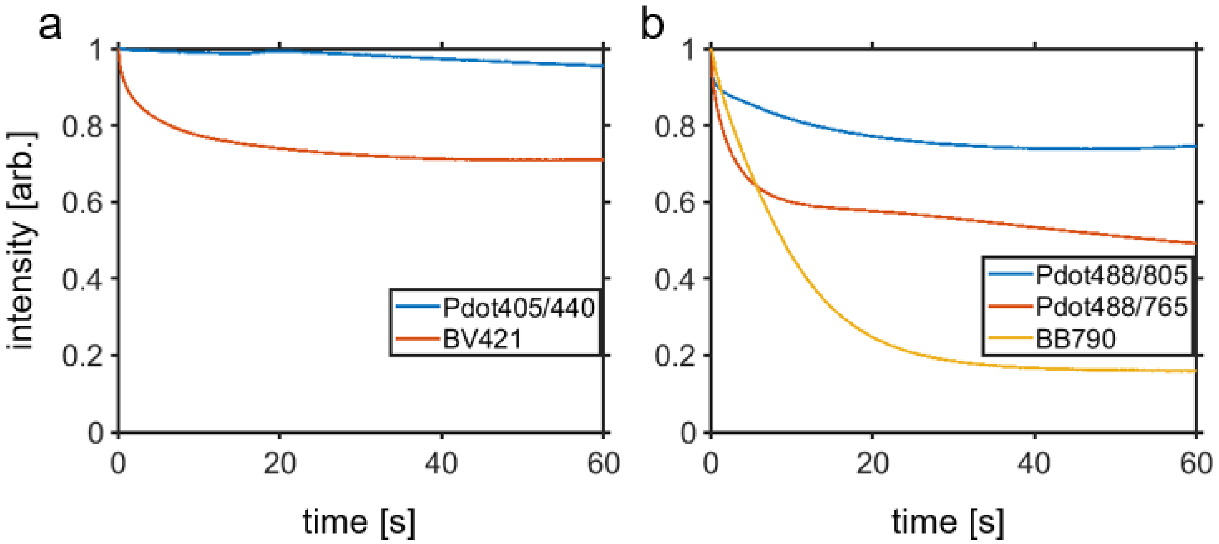


**Fig. S1. Comparing the photostability of Pdots against other dyes commonly used in flow cytometry.** (**a**) Comparison of the photostability of Pdot405/440 and Brilliant Violet 421, both excited at 405 nm. (**b**) Comparison of the photostability of the long Stokes-shift probes Pdot488/765, Pdot488/805, and BB790, all excited at 488 nm. All probes were illuminated at intensities comparable to that used for imaging.

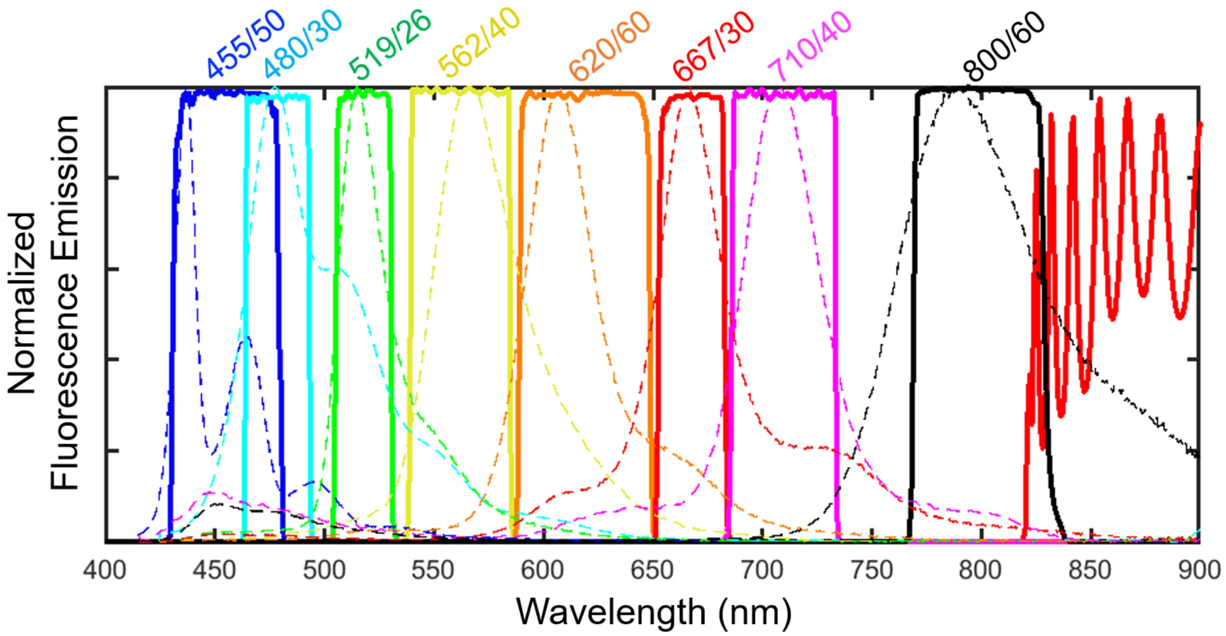


**Fig. S2. Emission spectra and associated filters overlay.** Emission spectra of eight 405-excited Pdots (dashed lines) along with the associated dichroic filter reflectance (red oscillating line above 825 nm) and emission filter transmission (solid colored lines).

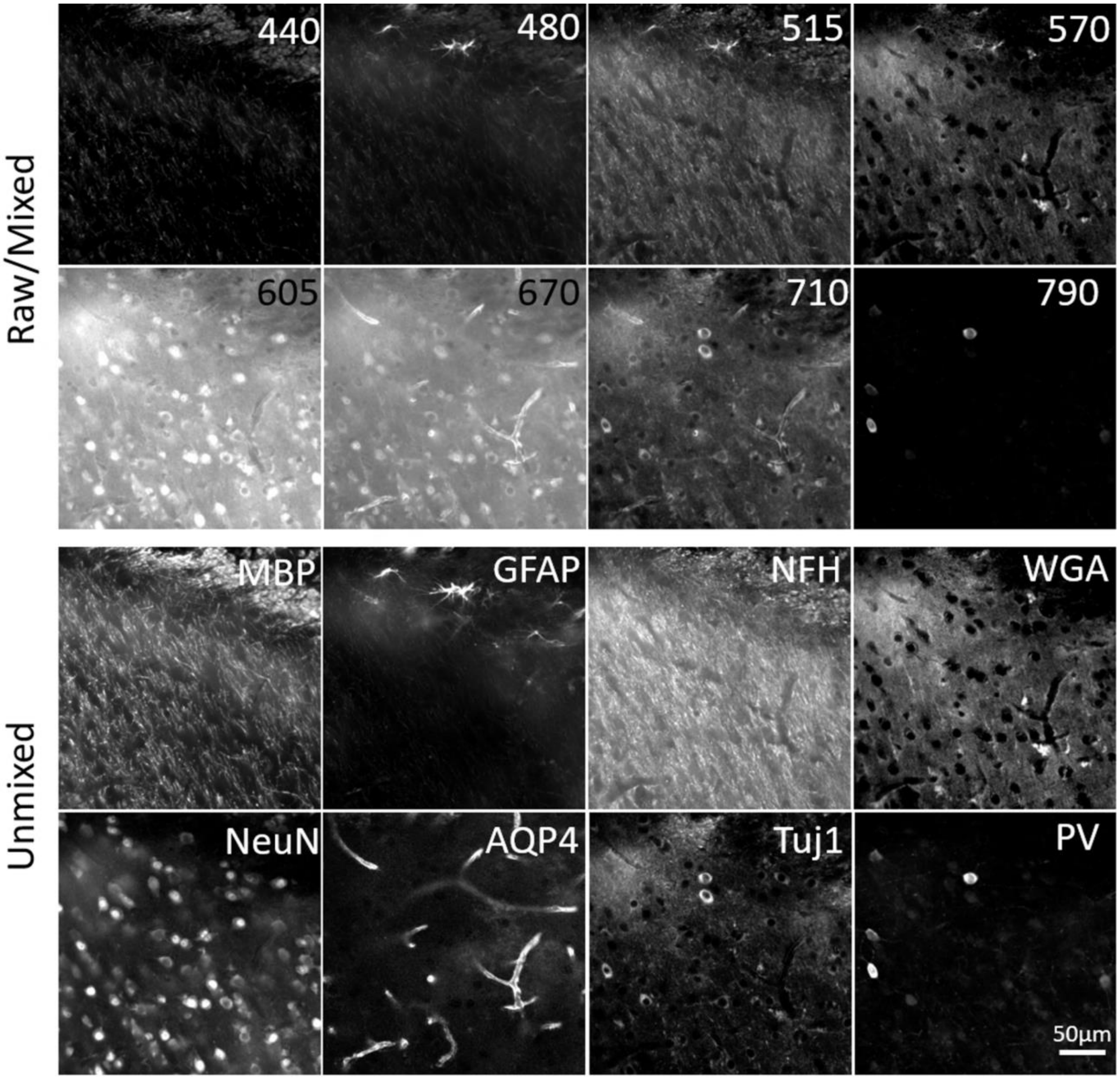


**Fig. S3. Raw and processed image comparison for the data set shown in Fig. 2.** Comparison of raw (mixed) images acquired at different emission wavelengths and the processed (unmixed) images that correspond to the primary antibody targets in mouse brain tissue.

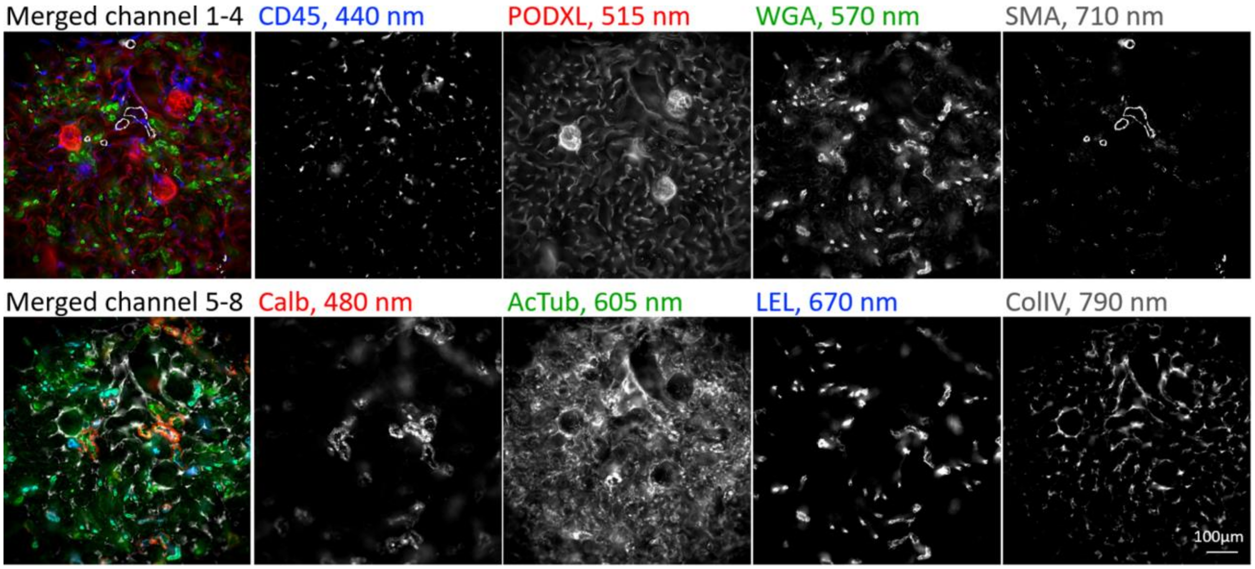


**Fig. S4. Eight-channel emission multiplexing on mouse kidney.** Unmixed images from indirect immunostaining of a 50 μm kidney slice using eight 405 nm-excited Pdot conjugates.

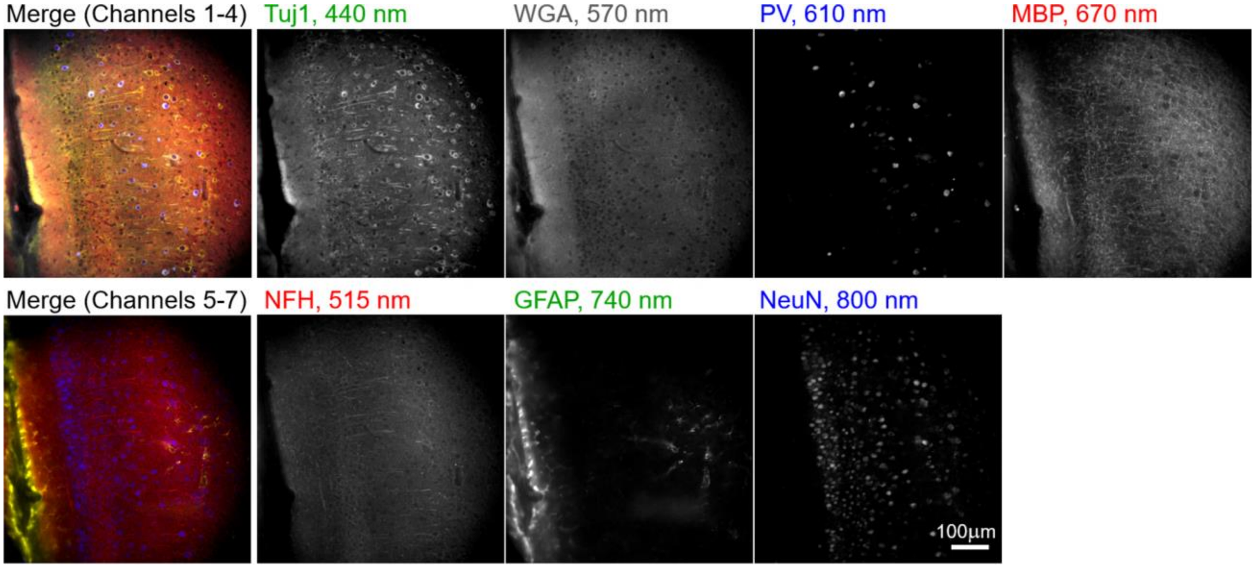


**Fig. S5. Seven-channel emission multiplexing on mouse brain.** Unmixed images from indirect immunofluorescence staining of a 100 μm thick mouse brain slice using seven 355 nm-excited Pdot conjugates.

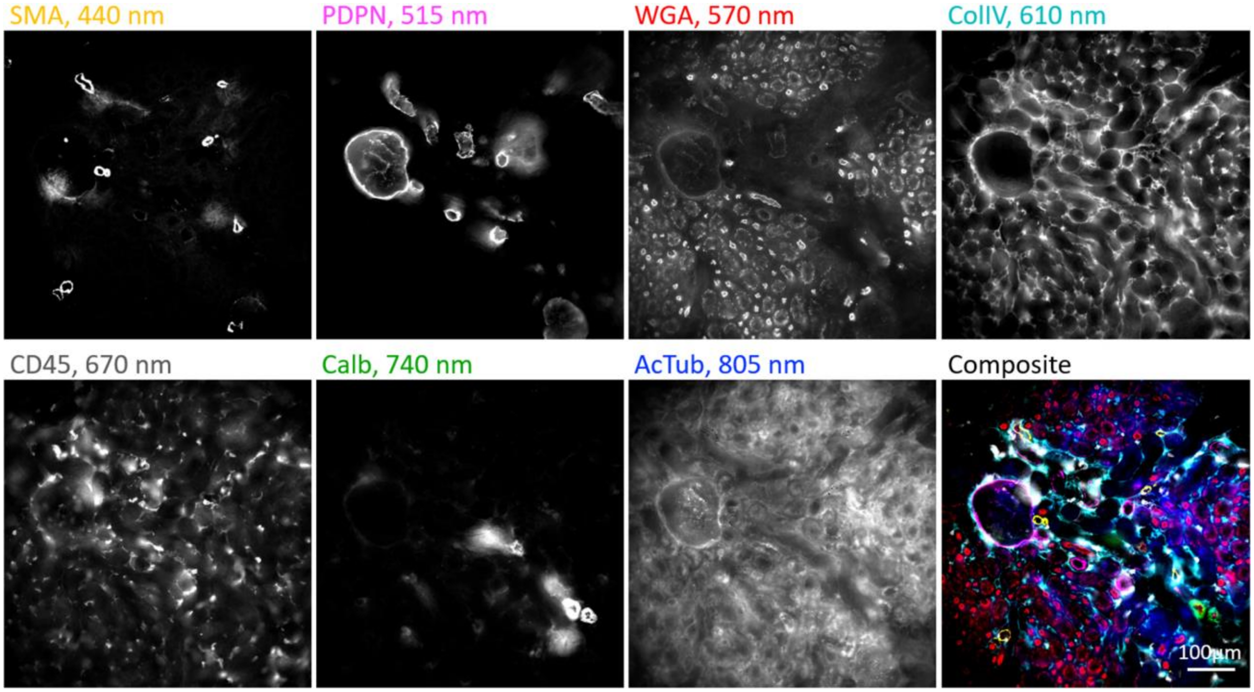


**Fig S6. Seven-channel emission multiplexing on mouse kidney.** Unmixed images from indirect immunostaining of a 50 μm kidney slice using seven 355 nm-excited Pdot conjugates.

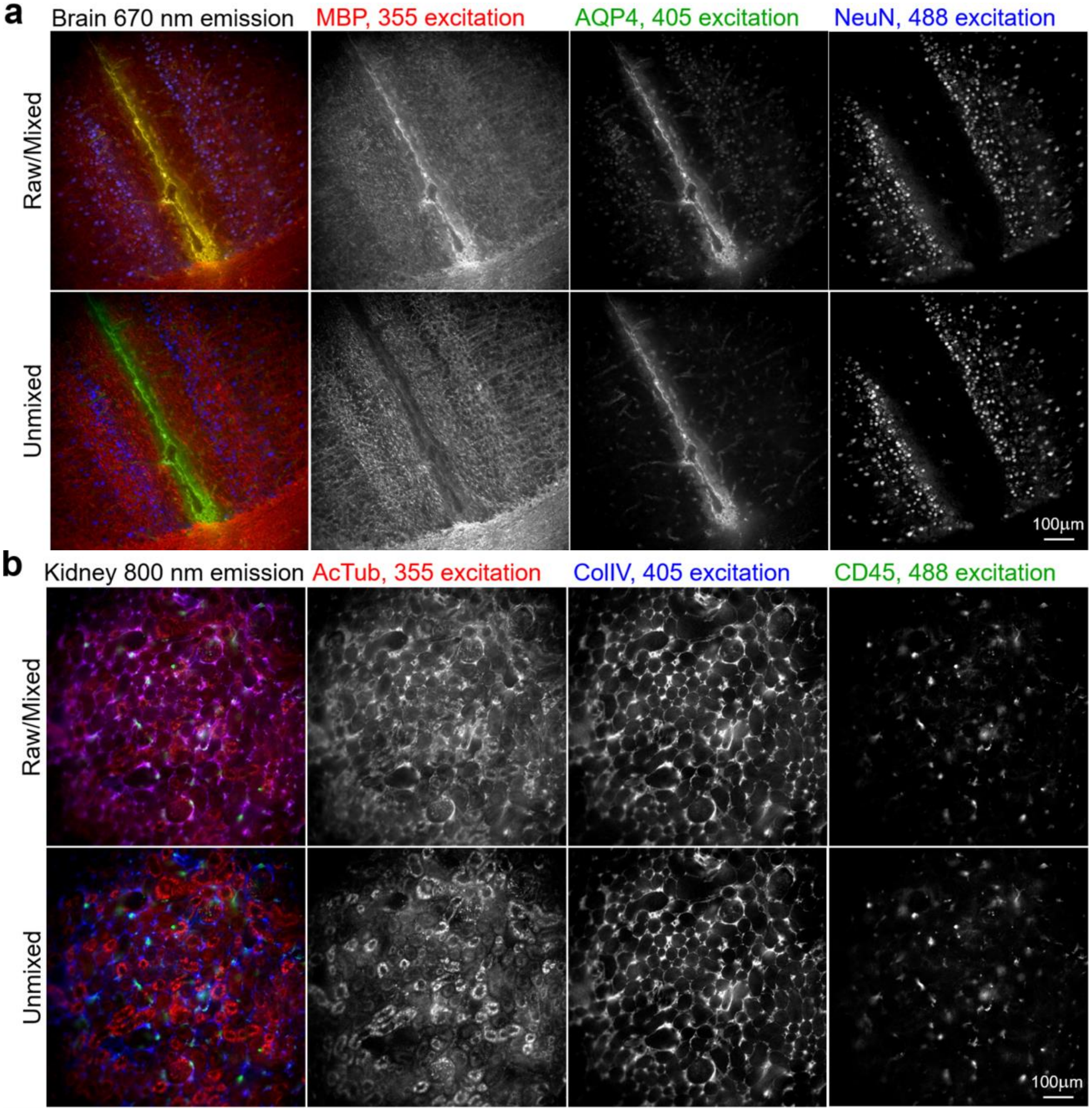


**Fig. S7. Three-channel excitation multiplexing on mouse brain and kidney.** A comparison of mixed (rows 1,3) and linearly unmixed (row 2,4) images for the data set shown in Fig. 3. (**a**): A 100 µm thick mouse brain slice was stained with three Pdot-antibody conjugates using three Pdots with different excitation maxima, and emission was collected in the same 670 nm channel. (**b**): A 100 µm thick mouse kidney slice was stained with three Pdot-antibody conjugates using three Pdots with different excitation maxima, and emission was collected in the same 800 nm channel.

**Table S1. Summary of sample labeling and imaging setup.** Other setup for all experiments (see method): used same microscope, exposure (100 ms), 20x objective lens, SOLA light engine, camera, immunostaining procedure, and incubation time; secondary antibodies are all from Jackson ImmunoResearch.

| **Fig.<br>Tissue<br>Thickness** | **Primary Antibody** | **Secondary Antibody** | **Filter Combinations** | | |
|---|---|---|---|---|---|
| 2, S3<br>Brain (Mouse 1)<br>100 µm section | MBP<br>GFAP<br>NFH<br>WGA<br>NeuN<br>AQP4<br>Tuj1<br>PV | Pdot405/440 Donkey anti-Rat<br>Pdot405/480 Donkey anti-Mouse IgG1<br>Pdot405/515 Goat anti-Chicken<br>Pdot405/570 WGA<br>Pdot405/605 Goat anti-Mouse IgG2b<br>Pdot405/670 Goat anti-Mouse IgG3<br>Pdot405/710 Goat anti-Mouse IgG2a<br>Pdot405/790 Donkey anti-Rabbit | *Dichroic:*<br>DI02-R405-25X36<br>Same for all | *Excitation filter:*<br>zet405/20x<br>Same for all | *Emission filters:*<br>ET455/50m<br>ET480/30m<br>ET519/26m<br>FF01-562/40<br>ET620/60m<br>ET667/30m<br>FF01-710/40<br>ET800/60 |
| 3A, S7<br>Brain (Mouse 1)<br>100 µm section | MBP<br>AQP4<br>NeuN | Pdot355/670 Donkey anti-Rat<br>Pdot405/670 Goat anti-Mouse IgG3<br>Pdot488/675 Goat anti-Mouse IgG2b | *Dichroics:*<br>FF376-DI01-25X36<br>DI02-R405-25X36<br>DI02-R488-25X36 | *Excitation filters:*<br>zet365/20x<br>zet405/20x<br>zet488/10x | *Emission filter:*<br>ET667/30m<br>ET667/30m<br>ET667/30m |
| 3B, S7<br>Kidney (Mouse 1)<br>100 µm section | AcTub<br>ColIV<br>CD45 | Pdot355/800 Goat anti-Mouse IgG2b<br>Pdot405/790 Donkey anti-Rabbit<br>Pdot488/805 Donkey anti-Rat | *Dichroics:*<br>FF376-DI01-25X36<br>DI02-R405-25X36<br>DI02-R488-25X36 | *Excitation filters:*<br>zet365/20x<br>zet405/20x<br>zet488/10x | *Emission filter:*<br>ET800/60<br>ET800/60<br>ET800/60 |
| 4<br>Brain (Mouse 3)<br>50 µm section | Pdot355/400 SOX2 (1)<br>Pdot355/515 LMNA/C (1)<br>Pdot355/570 MAP2<br>Pdot355/740 VEGFR3 (1)<br>Pdot405/440 GFAB<br>Pdot405/570 WGA<br>Pdot405/605 NeuN<br>Pdot405/670 TH<br>Pdot405/710 PV<br>Pdot405/790 MBP<br>Pdot488/515 Calb (1)<br>Pdot488/610 NFH<br>Pdot488/675 Tuj1<br>Pdot488/765 Nes | All primary conjugates, no secondaries necessary | *Dichroics:*<br>FF376-DI01-25X36<br>FF376-DI01-25X36<br>FF376-DI01-25X36<br>FF376-DI01-25X36<br>DI02-R405-25X36<br>DI02-R405-25X36<br>DI02-R405-25X36<br>DI02-R405-25X36<br>DI02-R405-25X36<br>DI02-R405-25X36<br>DI02-R488-25X36<br>DI02-R488-25X36<br>DI02-R488-25X36<br>DI02-R488-25X36 | *Excitation filters:*<br>zet365/20x<br>zet365/20x<br>zet365/20x<br>zet365/20x<br>zet405/20x<br>zet405/20x<br>zet405/20x<br>zet405/20x<br>zet405/20x<br>zet405/20x<br>zet488/10x<br>zet488/10x<br>zet488/10x<br>zet488/10x | *Emission filters:*<br>ET405/30m<br>ET519/26m<br>FF01-562/40<br>ET740/40x<br>ET455/50m<br>FF01-562/40<br>ET620/60m<br>FF01-562/40<br>FF01-710/40<br>ET800/60<br>ET519/26m<br>ET620/60m<br>ET667/30m<br>ET760/40 |
| 5<br>Kidney (Mouse 2)<br>50 µm section | Pdot355/440 PNA<br>Pdot355/515 CD13<br>Pdot355/570 AQP2<br>Pdot355/610 ITGA8<br>Pdot355/670 AQP1<br>Pdot355/740 VEGFR3 (1)<br>Pdot355/800 UMOD<br>Pdot405/440 CD45<br>Pdot405/480 Megalin<br>Pdot405/515 PODXL<br>Pdot405/570 WGA<br>Pdot405/605 ColIV<br>Pdot405/670 LEL<br>Pdot405/710 Agrin<br>Pdot405/790 α-SMA<br>Pdot488/515 Calb (1)<br>Pdot488/580 PDPL<br>Pdot488/610 REN<br>Pdot488/675 AcTub<br>Pdot488/765 Nes<br>Pdot488/805 PV | All primary conjugates, no secondaries necessary | *Dichroics:*<br>FF376-DI01-25X36<br>FF376-DI01-25X36<br>FF376-DI01-25X36<br>FF376-DI01-25X36<br>FF376-DI01-25X36<br>FF376-DI01-25X36<br>FF376-DI01-25X36<br>DI02-R405-25X36<br>DI02-R405-25X36<br>DI02-R405-25X36<br>DI02-R405-25X36<br>DI02-R405-25X36<br>DI02-R405-25X36<br>DI02-R405-25X36<br>DI02-R405-25X36<br>DI02-R488-25X36<br>DI02-R488-25X36<br>DI02-R488-25X36<br>DI02-R488-25X36<br>DI02-R488-25X36<br>DI02-R488-25X36 | *Excitation filters:*<br>zet365/20x<br>zet365/20x<br>zet365/20x<br>zet365/20x<br>zet365/20x<br>zet365/20x<br>zet365/20x<br>zet405/20x<br>zet405/20x<br>zet405/20x<br>zet405/20x<br>zet405/20x<br>zet405/20x<br>zet405/20x<br>zet405/20x<br>zet488/10x<br>zet488/10x<br>zet488/10x<br>zet488/10x<br>zet488/10x<br>zet488/10x | *Emission filters:*<br>ET455/50m<br>ET519/26m<br>FF01-562/40<br>ET620/60m<br>ET667/30m<br>ET740/40x<br>ET800/60<br>ET455/50m<br>ET480/30m<br>ET519/26m<br>FF01-562/40<br>ET620/60m<br>ET667/30m<br>FF01-710/40<br>ET800/60<br>ET519/26m<br>FF01-562/40<br>ET620/60m<br>ET667/30m<br>ET760/40<br>ET800/60 |
| S4<br>Kidney (Mouse 2)<br>50 µm section | CD45<br>Calb<br>PODXL<br>WGA<br>AcTub<br>LEL<br>α-SMA<br>ColIV | Pdot405/440 Donkey anti-Rat<br>Pdot405/480 Donkey anti-Mouse IgG1<br>Pdot405/515 PODXL<br>Pdot405/570 WGA<br>Pdot405/605 Goat anti-Mouse IgG2b<br>Pdot405/670 LEL<br>Pdot405/710 Goat anti-Mouse IgG2a<br>Pdot405/790 Donkey anti-Rabbit | *Dichroic:* DI02-R405-25X36<br>Same for all | *Excitation filter:*<br>zet405/20x<br>Same for all | *Emission filters:*<br>ET455/50m<br>ET480/30m<br>ET519/26m<br>FF01-562/40<br>ET620/60m<br>ET667/30m<br>FF01-710/40<br>ET800/60 |

| | | | | | |
|---|---|---|---|---|---|
| S5<br>Brain (Mouse 1)<br>100 μm section | Tuj1<br>NFH<br>WGA<br>PV<br>MBP<br>GFAB<br>NeuN | Pdot355/440 Goat anti-mouse IgG2a<br>Pdot355/515 Donkey anti-Chicken<br>Pdot355/570 WGA<br>Pdot355/610 Donkey anti-Rabbit<br>Pdot355/670 Donkey anti-Rat<br>Pdot355/740 Goat anti-Mouse IgG1<br>Pdot355/800 Goat anti-Mouse IgG2b | *Dichroics:*<br>FF376-DI01-25X36<br>Same for all | *Excitation filters:*<br>zet365/20x<br>Same for all | *Emission filters:*<br>ET455/50m<br>ET519/26m<br>FF01-562/40<br>ET620/60m<br>ET667/30m<br>ET740/40x<br>ET800/60 |
| S6<br>Kidney (Mouse 2)<br>50 μm section | α-SMA<br>PDPN<br>WGA<br>ColIV<br>CD45<br>Calb<br>AcTub | Pdot355/440 Goat anti-mouse IgG2a<br>Pdot355/515 Donkey anti-Hamster<br>Pdot355/570 WGA<br>Pdot355/610 Donkey anti-Rabbit<br>Pdot355/670 Donkey anti-Rat<br>Pdot355/740 Goat anti-Mouse IgG1<br>Pdot355/800 Goat anti-Mouse IgG2b | *Dichroics:*<br>FF376-DI01-25X36<br>Same for all | *Excitation filters:*<br>zet365/20x<br>Same for all | *Emission filters:*<br>ET455/50m<br>ET519/26m<br>FF01-562/40<br>ET620/60m<br>ET667/30m<br>ET740/40x<br>ET800/60 |

### Table S2. Pdot-secondary antibody conjugates.

| | Primary Antibody or Lectin | Protein Target or Lectin | Host | Vendor | Catalog # | Clonality | Pdot Conjugated Secondary Antibody or Lectin | Catalog # |
|---|---|---|---|---|---|---|---|---|
| Brain - 355ex | Tuj1/TUBB3 | Tubulin β 3 | Mouse IgG2a | BioLegend | 801201 | monoclonal | Pdot355/440 Goat anti-mouse IgG2a | 115-005-206 |
| | NFH | Neurofilament Heavy Chain | Chicken | Aves Labs | NFH | | Pdot355/515 Donkey anti-Chicken | 703-005-155 |
| | WGA | Wheat Germ Agglutinin | | Vector Labs | L-1020 | | Pdot355/570 WGA | - |
| | PV | Parvalbumin | Rabbit | Abcam | ab11427 | polyclonal | Pdot355/610 Donkey anti-Rabbit | 711-005-152 |
| | MBP | Myelin Basic Protein | Rat | Abcam | ab7349 | monoclonal | Pdot355/670 Donkey anti-Rat | 715-005-151 |
| | GFAP | Glial Fibrillary Acidic Protein | Mouse IgG1 | Cell Signaling | 3670S | monoclonal | Pdot355/740 Goat anti-mouse IgG1 | 115-005-205 |
| | NeuN | Neuronal Nuclei | Mouse IgG2b | Encor | MCA-1B7 | monoclonal | Pdot355/800 Goat anti-mouse IgG2b | 115-005-207 |
| Kidney - 355ex | α-SMA | alpha-Smooth Muscle Actin | Mouse IgG2a | BioLegend | 904601 | monoclonal (1A4) | Pdot355/440 Goat anti-mouse IgG2a | 115-005-206 |
| | PDPN | Podoplanin | Syrian Hamster IgG | DSHB | 8.1.1-s | monoclonal | Pdot355/515 Donkey anti-Hamster | Unavailable |
| | WGA | Wheat Germ Agglutinin | | Vector Labs | L-1020 | | Pdot355/570 WGA | - |
| | AQP1 | Aquaporin 1 | Rabbit | Abcam | ab15080 | polyclonal | Pdot355/610 Donkey anti-Rabbit | 711-005-152 |
| | CD45 | Leukocyte Common Antigen | Rat | BioLegend | 103101 | monoclonal (30-F11) | Pdot355/670 Donkey anti-Rat | 715-005-151 |
| | Calb | Calbindin | Mouse IgG1 | Abcam | ab82812 | monoclonal | Pdot355/740 Goat anti-mouse IgG1 | 115-005-205 |
| | AcTub | Acetylated Tubulin | Mouse IgG2b | Sigma | T7451-25UL | monoclonal (6-11B-1) | Pdot355/800 Goat anti-mouse IgG2b | 115-005-207 |
| Brain - 405ex | MBP | Myelin Basic Protein | Rat | Abcam | ab7349 | monoclonal | Pdot405/440 Donkey anti-Rat | 715-005-151 |
| | GFAP | Glial fibrillary acidic protein | Mouse IgG1 | Cell Signaling | 3670S | monoclonal | Pdot405/480 Donkey anti-mouse IgG1 | 115-005-205 |
| | NFH | Neurofilament Heavy-chain | Chicken | Aves Labs | NFH | polyclonal | Pdot405/515 Goat anti-chicken | 103-005-155 |
| | WGA | Wheat Germ Agglutinin | | Vector Labs | L-1020 | | Pdot405/570 WGA | - |
| | NeuN | Neuronal Nuclei | Mouse IgG2b | Encor | MCA-1B7 | monoclonal | Pdot405/605 Goat anti-mouse IgG2b | 115-005-207 |
| | AQP4 | Aquaporin 4 | Mouse IgG3 | Santa Cruz Biotechnology | sc-32739 | monoclonal | Pdot405/670 Goat anti-mouse IgG3 | 115-005-209 |
| | Tuj1/TUBB3 | Tubulin β 3 | Mouse IgG2a | BioLegend | 801201 | monoclonal | Pdot405/710 Goat anti-mouse IgG2a | 115-005-206 |
| | PV | Parvalbumin | Rabbit | Abcam | ab11427 | polyclonal | Pdot405/790 Donkey anti-rabbit | 711-005-152 |
| Kidney - 405ex | CD45 | Leukocyte Common Antigen | Rat IgG2b | BioLegend | 103101 | monoclonal (30-F11) | Pdot405/440 Donkey anti-Rat | 715-005-151 |
| | Calb | Calbindin | Mouse IgG1 | Abcam | ab82812 | monoclonal | Pdot405/480 Donkey anti-mouse IgG1 | 115-005-205 |
| | PODXL | Podocalyxin | Goat | R&D Systems | AF1556 | | Pdot405/515 Goat anti-PODXL | - |
| | WGA | Wheat Germ Agglutinin | | Vector Labs | L-1020 | | Pdot405/570 WGA | - |
| | AcTub | Acetylated Tubulin | Mouse IgG2b | Sigma | T7451-25UL | monoclonal (6-11B-1) | Pdot405/605 Goat anti-mouse IgG2b | 115-005-207 |
| | LEL | Lycopersicon Esculentum Tomato Lectin | | Vector Labs | L-1170-2 | | Pdot405/670 LEL | - |
| | α-SMA | alpha-Smooth Muscle Actin | Mouse IgG2a | BioLegend | 904601 | monoclonal (1A4) | Pdot405/710 Goat anti-mouse IgG2a | 115-005-206 |
| | ColIV | Collagen IV | Rabbit | Abcam | ab6586 | polyclonal | Pdot405/790 Donkey anti-rabbit | 711-005-152 |
| Brain - 488ex | Tuj1/TUBB3 | Tubulin β 3 | Mouse IgG2a | BioLegend | 801201 | monoclonal | Pdot488/610-Goat anti-mouse IgG2a | 115-005-206 |
| | NeuN | Neuronal Nuclei | Mouse IgG2b | Encor | MCA-1B7 | monoclonal | Pdot488/675-Goat anti-mouse IgG2b | 115-005-207 |
| | MBP | Myelin Basic Protein | Rat | Abcam | ab7349 | monoclonal | Pdot488/805-Donkey anti-Rat | 715-005-151 |
| Kidney - 488ex | α-SMA | alpha-Smooth Muscle Actin | Mouse IgG2a | BioLegend | 904601 | monoclonal (1A4) | Pdot488/610-Goat anti-mouse IgG2a | 115-005-206 |
| | AcTub | Acetylated Tubulin | Mouse IgG2b | Sigma | T7451-25UL | monoclonal (6-11B-1) | Pdot488/675-Goat anti-mouse IgG2b | 115-005-207 |
| | CD45 | Leukocyte Common Antigen | Rat | BioLegend | 103101 | monoclonal (30-F11) | Pdot488/805-Donkey anti-Rat | 715-005-151 |

### Table S3. Pdot-primary antibody conjugates.

| Pdot | Antibody or Lectin | Protein Target or Lectin | Performance | Host | Vendor | Catalog # |
|---|---|---|---|---|---|---|
| 355-400 | CD31/PECAM-1 | Cluster of Designation 31, Platelet Endothelial Cell Adhesion Molecule-1 | ++ | Goat | R&D Systems | AF3628 |
| 355-400* | SOX2 (1) | Sex Determining Region Y-Box 2 | +++ | Goat | Novus Biologicals | AF2018 |
| 355-400 | SOX2 (2) | Sex Determining Region Y-Box 2 | ++ | Mouse IgG1 | Thermo Fisher | 66411-1-IG |
| 355-440 | Tuj1/TUBB3 | Tubulin β 3 | + | Mouse IgG2a | BioLegend | 801201 |
| 355-440† | PNA | Peanut Agglutinin | +++ | - | Vector | L-1070-25 |
| 355-440 | SYNPO | Synaptopodin | + | Rabbit | Thermo Fisher | TA890150 |
| 355-515† | CD13/APN | Cluster of Designation 13, Aminopeptidase N | +++ | Rat IgG2a | Thermo Fisher | MA1-33449 |
| 355-515* | LMNA/C | Lamin A/C | +++ | Rabbit | Proteintech | 10298-1-AP |
| 355-515 | Podocin (1) | Podocin | + | Rabbit | Millipore Sigma | P0372-200UL |
| 355-570† | AQP2 | Aquaporin 2 | +++ | Rabbit | Novus Biological | NB110-74682 |
| 355-570* | MAP2 | Microtubule Associated Protein 2 | +++ | Chicken IgGY | Novus Biologicals | NB300-213 |
| 355-610 | AQP4 (1) | Aquaporin-4 | + | Rabbit | Millipore Sigma | HPA014784-100UL |
| 355-610 | AQP4 (2) | Aquaporin-4 | + | Rabbit | LSBio | LS-C150456 |
| 355-610 | Nephrin | Nephrin | + | Goat | R&D Systems | AF3159 |
| 355-610 | CD45 | Cluster of Designation 45, Leukocyte Common Antigen | +++ | Rat IgG2b | BioLegend | 103102 |
| 355-610† | ITGA8 | Integrin alpha 8 | +++ | Goat | Novus Biologicals | AF4076 |
| 355-610 | Podocin (2) | Podocin | + | Rabbit | Thermo Fisher | PA5-37284 |
| 355-670† | AQP1 | Aquaporin 1 | +++ | Rabbit | Proteintech | 20333-1-AP |
| 355-670 | PNA | Peanut Agglutinin | ++ | - | Vector | L-1070-5 |
| 355-740*† | VEGFR3 | Vascular Endothelial Growth Factor Receptor 3 | +++ | Rat | Novus Biologicals | NB110-61018, 2022 |
| 355-740 | VIM | Vimentin | ++ | IgG2a | R&D Systems | MAB21052-100 |
| 355-805 | GAD67 | Glutamate Decarboxylase 1 | + | Rabbit | LSBio | LS-C827246-100 |
| 355-805 | CD90/Thy1 | Cluster of Designation 90, Thymus Cell Antigen-1 | + | Rat IgG1 | Novus Biologicals | NB100-65543 |
| 355-805† | UMOD/THP | Uromodulin, Tamm-Horsfall Protein | +++ | Rat IgG2a | R&D Systems | MAB5175 |
| 405-440† | CD45 | Cluster of Designation 45, Leukocyte Common Antigen | +++ | Rat IgG2b | BioLegend | 103102 |
| 405-440* | GFAP | Glial Fibrillary Acidic Protein | +++ | Chicken IgY | Thermo Fisher | PA1-10004 |
| 405-480 | CD31/PECAM-1 | Cluster of Designation 45, Platelet Endothelial Cell Adhesion Molecule-1 | ++ | Goat | R&D Systems | AF3628 |
| 405-480† | Megalin/LRP2 | Low Density Lipoprotein-related Protein 2 | +++ | Rabbit | Invitrogen | PA5-92032 |
| 405-515 | NFL | Neurofilament Light Chain | + | Chicken | Aves Labs | NFL |
| 405-515† | PODXL | Podocalyxin | +++ | Goat | R&D Systems | AF1556 |
| 405-570*† | WGA | Wheat Germ Agglutinin | +++ | - | Vector | L-1020-25 |
| 405-610† | ColIV | Collagen IV | +++ | Rabbit | Abcam | ab6586 |
| 405-605* | NeuN | Neuronal Nuclei | +++ | Mouse IgG2b | BioLegend | 834501 |
| 405-670† | LEL | Lycopersicon Esculentum Tomato Lectin | +++ | - | Vector | L-1170-2 |
| 405-670* | TH | Tyrosine Hydroxylase | ++ | Mouse IgG2a | BioLegend | 818001 |
| 405-710† | Agrin | Agrin | +++ | Goat | R&D Systems | AF550 |
| 405-710* | PV | Parvalbumin | +++ | Rabbit | Novus Biologicals | NB120-11427 |
| 405-790† | α-SMA | alpha-Smooth Muscle Actin | +++ | Mouse IgG2a | Millipore Sigma | A5228-200UL |
| 405-790* | MBP | Myelin Basic Protein | +++ | Mouse IgG2b | Invitrogen | MA1-10837 |
| 488-520*† | Calb (1) | Calbindin | +++ | Rabbit | LSBio | LS-B16562-100 |
| 488-520 | Calb (2) | Calbindin | ++ | Mouse IgG1 | Abcam | ab82812 |
| 488-520 | Calb (3) | Calbindin | + | Chicken | Novus Biologicals | NBP2-50028 |
| 488-580† | PDPN | Podoplanin | ++ | Goat | R&D Systems | AF3244 |
| 488-580 | LEL | Lycopersicon Esculentum Tomato Lectin | +++ | - | Vector | L-1170-2 |
| 488-610* | NFH | Neurofilament Heavy Chain | +++ | Chicken IgY | Aves Labs | NFH |

| | | | | | | |
|---|---|---|---|---|---|---|
| 488-610† | REN | Renin | +++ | Goat | R&D Systems | AF4277 |
| 488-675† | AcTub | Acetylated Tubulin | +++ | Mouse IgG2b | Millipore Sigma | T7451-200UL |
| 488-675* | Tuj1/TUBB3 | Tubulin β 3 | +++ | Mouse IgG2b | BioLegend | MMS-435P |
| 488-765*† | Nes/Nestin | Neuroepithelial stem cell protein | +++ | Mouse IgG2a | R&D Systems | MAB2736 |
| 488-765 | SYNPO (1) | Synaptopodin | + | Rabbit | Millipore Sigma | S9442-200UL |
| 488-810 | CK8 | Cytokeratin 8 | + | Rabbit | LSBio | LS-B7928-50 |
| 488-810 | LMNB1 (1) | Lamin B1 | ++ | Mouse IgG1 | Proteintech | 66095-1-Ig |
| 488-810† | PV | Parvalbumin | +++ | Rabbit | Novus Biologicals | NB120-11427 |

* labeled are used in brain 14-panel stain
† labeled are used in kidney 21-panel stain
+++ Positive and correct signals in multiple samples
++ Positive and correct signals in some tested samples
+ Images dominated by autofluorescence or exhibiting incorrect non-specific signals in most samples

### Table S4. Optical filters.

| | Chroma | Semrock |
|---|---|---|
| **Dichroics (longpass)** | | DI02-R405-25X36 |
| | | DI02-R488-25X36 |
| | | FF376-DI01-25X36 |
| **Excitation filters** | zet365/20x | |
| | zet405/20x | |
| | zet488/10x | |
| **Emission filters** | ET405/30m | FF01-562/40 |
| | ET455/50m | FF01-710/40 |
| | ET480/30m | |
| | ET519/26m | |
| | ET620/60m | |
| | ET667/30m | |
| | ET740/40x | |
| | ET760/40 | |
| | ET800/60 | |
